# Neuronal Classification from Network Connectivity via Adjacency Spectral Embedding

**DOI:** 10.1101/2020.06.18.160259

**Authors:** Ketan Mehta, Rebecca F. Goldin, David Marchette, Joshua T. Vogelstein, Carey E. Priebe, Giorgio A. Ascoli

## Abstract

This work presents a novel strategy for classifying neurons, represented by nodes of a directed graph, based on their circuitry (edge connectivity). We assume a stochastic block model (SBM) where neurons belong together if they connect to neurons of other groups according to the same probability distributions. Following adjacency spectral embedding (ASE) of the SBM graph, we derive the number of classes and assign each neuron to a class with a Gaussian mixture model-based expectation-maximization (EM) clustering algorithm. To improve accuracy, we introduce a simple variation using random hierarchical agglomerative clustering to initialize the EM algorithm and picking the best solution over multiple EM restarts. We test this procedure on a large (*n* ~ 2^12^ − 2^15^ neurons), sparse, biologically inspired connectome with eight neuron classes. The simulation results demonstrate that the proposed approach is broadly stable to the choice of dimensional embedding and scales extremely well as the number of neurons in the network increases. Clustering accuracy is robust to variations in model parameters and highly tolerant to simulated experimental noise, achieving perfect classifications with up to 40% of swapped edges. Thus, this approach may be useful to analyze and interpret large-scale brain connectomics data in terms of underlying cellular components.

## 1 INTRODUCTION

A functionally relevant, quantitative description of cellular diversity in the brain remains a pressing open problem in neuroscience. Traditionally, investigators have classified neurons by subsets of multifarious properties, including physiology, biochemistry, and morphology (e.g. a fast-spiking, parvalbumin-expressing, aspiny interneuron). In spite of the widespread and foundational use of the notion of cell class, there is no formal definition of this concept, and how exactly a cell class relates to network connectivity remains a matter of considerable debate in the community (Ascoli et al. 2008; DeFelipe et al. 2013). In particular, given a “solved” connectome (a complete list of all neurons and their connections), is it possible to objectively find the number of neuronal connectivity classes, and to assign each neuron to a class? This would also answer the related open question of how many cell classes there are from the connectomics perspective.

In this work we introduce a novel strategy for classifying neurons based on their circuitry. In particular, after formalizing the concept of cell class based on network connectivity, we present a technique to derive the number of cell classes from a neuronal connectome, and to assign each neuron to a class. Using neurobiologically realistic surrogate data, we demonstrate that this technique is robust and efficient.

We begin by asking a mathematical question derived from the neuroscientific one. Recall that a **directed graph** (*V, E*) consists of vertices *V* (a finite set), and directed edges *E*, a subset of ordered pairs of *V × V*. We assume the directed graph is **simple**, i.e., there is at most one edge between any two distinct vertices, and no edge from a vertex to itself, though we allow the possibility of edges in either direction. For the purpose of our analysis, each connectome may be represented by such a directed graph, wherein the vertex represents a neuron and the edge represents a directed synaptic (usually axon-dendrite) connection. Further, we adopt a generative model approach by using a stochastic block model (SBM) to add additional structure to the directed graph. In this model vertices are partitioned into nonoverlapping groups called blocks, such that the probability of an edge between two vertices depends only on their respective block memberships. Vertices in the same block are thus stochastically equivalent. Given a directed SBM graph, our goal is then to estimate the number of blocks and assign each vertex to its respective block.

Recently, SBMs have been successfully used to model connectomes (Moyer et al. 2015; Pavlovic et al. 2014), as well as to identify network community structures within connectomes (Betzel et al. 2018; Faskowitz et al. 2018). Our approach here, however, is different from these studies in two important aspects. First, we use surrogate connectomic data loosely inspired by the entorhinal-CA1 circuit of the rodent hippocampal formation. The scale and structure of the neuronal network analyzed in this work is therefore vastly different, with substantially larger graphs (~ 2^12^ − 2^15^ vertices) and sparse (~ 4%) connectivity. Second, and more fundamentally, our focus is on developing a robust mathematical framework using spectral graph clustering to capture the latent block structure of the directed graph. We are motivated by recent results (Sussman et al. 2012; Priebe et al. 2017, 2019) which demonstrate the use of adjacency spectral embedding (ASE) in conjunction with Gaussian mixture model (GMM)-based clustering to estimate block membership. Here we adopt and modify the GMM○ASE framework, and present a strategy to cluster large, sparse graphs modeled from surrogate connectomic data.

Given a graph, we begin by embedding it into a much lower dimensional space by computing the singular value decomposition of a slightly modified version of the adjacency matrix. Since we consider directed graphs, we embed a concatenation of the left and right singular vectors, which correspond to the outgoing (pre-synaptic) and incoming (post-synaptic) connections, respectively. Following the embedding, the latent vectors are modeled as a GMM and clustered using the expectation maximization (EM) algorithm. However, the convergence of the EM algorithm is highly sensitive to the starting values chosen to initialize the algorithm, especially for the multivariate GMM case (Biernacki et al. 2003; Kwedlo 2015; Shireman et al. 2017), and often gets trapped in a local optimum. Therefore, we propose using a multiple restart approach wherein we apply hierarchical agglomerative clustering to randomly initialize and start the EM algorithm multiple times, and subsequently pick the model with the largest value of Bayesian information criterion (BIC) over multiple restarts.

We perform a series of experimental simulations with surrogate data to validate the effectiveness of the proposed multiple random restart EM. The simulation results demonstrate the proposed clustering strategy to be extremely effective in successfully recovering the true number of classes and individual class assignment of the vertices. The random multiple restart approach also heavily outperforms GMM-based hierarchical partition initialization (Scrucca and Raftery 2015), while having the advantage of being broadly stable over a wide selection of embedding dimensions, as choosing an optimal value for dimensional embedding remains an open problem with spectral graph clustering in general. The proposed approach is also robust to variations in model parameters and scales extremely well as the number of neurons in the network increases. Moreover, our analysis shows this method to be highly tolerant to noise in the form of edge swaps akin to experimental errors in pre- or post-synaptic neuron identification.

The rest of the paper is organized as follows. We begin by describing our connectome model in Section 2. The adjacency spectral embedding framework in presented in Section 3, followed by the spectral graph clustering and the multiple random restart EM algorithm in Section 4. In Section 5 we present the simulation results. We finally point to the implications of this work to neuroscience in Section 6.

## 2 MODELING THE CONNECTOME

### 2.1 Stochastic Block Models

Consider a directed graph (*V, E*) that consists of vertices *V* (a finite set), and directed edges E, a subset of ordered pairs of *V × V*. We write (*υ,w*) ∈ E for *υ, w ∈ V* if there is a directed edge from *υ* to *w*. Further, we assume the graph to be **simple**, i.e., (*υ,w*) ∈ *E* implies *υ = w*. As *E* is a set of ordered pairs, there is at most one directed edge from any vertex *υ* to a distinct vertex *w*. We allow the possibility of edges (*υ, w*) and (*w, v*). We formally define a partitioned directed graph as follows:

#### Definition 2.1.

***A partitioned directed graph** is a triple* (*V, E, τ*), *where* (*V, E*) *is a simple directed graph and τ*: *V* → {1,…,*k*} *is an assignment of a number from 1 to k for each vertex in V*.

The map *τ* is also called a **block-assignment function**. For any such *τ*, we notate the subsets formed by the partition by

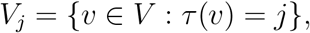

for *j* = 1,…, *k*. We call *j = τ*(*υ*) the **class** of *υ*. In a stochastic block model (SBM) (Holland and Leinhardt 1981; Holland et al. 1983), stochastically equivalent vertices are partitioned together into classes. In particular, SBMs make the assumption that there is a probability distribution with parameters associated with each ordered pair (*V_i_, V_j_*), such that vertices from the *i*th class connect to those in the *j*th class according to a specified random process, in our case a Bernoulli trial with parameter *p_i_j*. Let *P* = (*p_ij_*) be a matrix collecting these parameters.

We formally define the generative model of the standard directed SBM as follows.

#### Definition 2.2.

***A directed stochastic block model** (SBM) is a generative model for directed graphs. Let *n* be the number of nodes (vertices), k the number of groups (classes), P* = (*p_ij_*) ∈ [0,1]^*k×k*^ *the block connectivity probability matrix (edge probabilities), and τ*: *V* → {1,…,*k*} *the assignment of each node to a group. A directed SBM graph is a partitioned directed graph G* = (*V, E, τ*) *whose edges are independent Bernoulli draws with probability P*{(*υ, w*) ∈ *E*} = *p*_*τ*(*υ*),*τ*(*w*)_.

Let *ρ_j_*: = |*V_j_*|/*n* be the proportion of vertices in the *j*th group. The *k*-tuple *ρ*: = (*ρ*_1_,…, *ρ_k_*) indicates the proportional sizes of these classes. Note that {*V*_1_,…, *V_k_*} and *ρ* depend only on τ.

In a *general* SBM^1^ (Abbe 2018), the vertex assignment, and thus the class size |*V_j_*| of the generated graph, is subject to a random process, however in our generative model the assignment is instead specified by the block-assignment function τ.

### 2.2 Connectome Generation

The experimental design begins with using a directed SBM to generate stochastic realizations (simulations) of the biological connectome. The surrogate model used is loosely inspired by the entorhinal-CA1 circuit of the rodent hippocampal formation based on Hippocampome.org data (Wheeler et al. 2015). Specifically, we consider a directed neuronal network consisting of *n* cells, where *n* varies, and *k* = 8 distinct cell types. Each cell type is briefly described in Table 1. The model is parametrized by the connectivity probability matrix

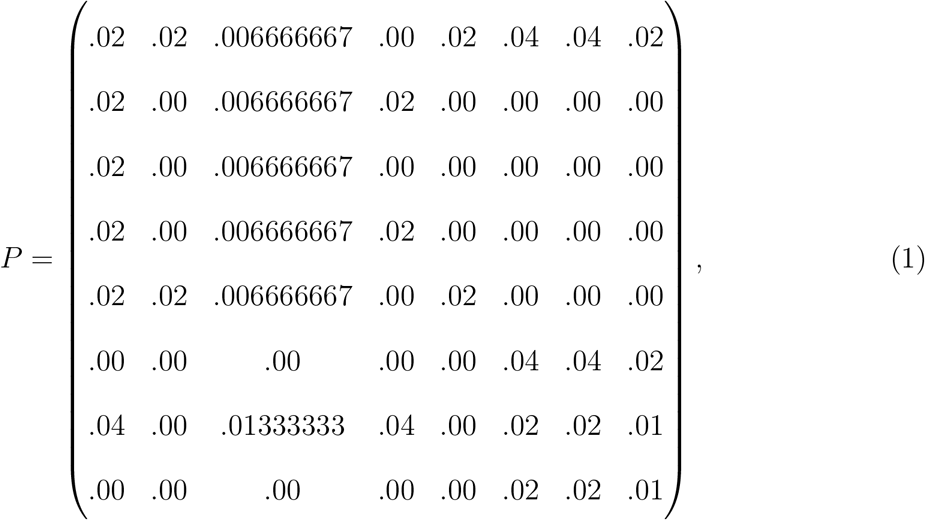

and a block membership vector *ρ* which denotes the proportions of the cells (vertices) assigned each cell type (class),

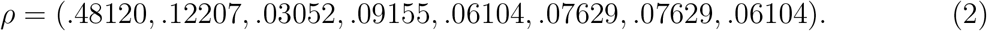

**Table 1:** The eight cell classes.

|  |  |
| --- | --- |
| CA1 Pyramidal | Principal output neurons of the hippocampus. One of the most studied and best characterized excitatory neurons of the mammalian brain. |
| CA1 Oriens/Lacunosum-Moleculare | Local inhibitory neurons. Dendrites are in the oriens layer and axons start in the oriens and go up to lacunosum-moleculare. |
| CA1 Basket | Local peri-somatic inhibitory interneurons. Axons target pyramidal and basket cells. Dendrites span all layers of CA1. |
| CA1 Perforant Pathway-Associated | Local inhibitory interneurons with axons and dendrites confined to the lacunosum-moleculare layer. |
| CA1 Oriens | Local inhibitory interneurons with dendrites and axons confined to the oriens layer. |
| Entorhinal Cortex Layer 5 Pyramidal | Deep layer excitatory neurons with dendrites and axons extending through the deep and superficial layers of the entorhinal cortex. |
| Entorhinal Cortex Layer 3 Pyramidal | Superficial layer excitatory neurons. Dendrites span through the deep and superficial layers of the entorhinal cortex; axons start in layer 3 and project to CA1 lacunosum-moleculare. |
| Entorhinal Cortex GABAergic Cells | Inhibitory local interneurons with axons and dendrites through the deep and superficial layers of the entorhinal cortex. |

The assignment *τ* of cells to cell types simply maps the first *nρ*_1_ cells to the first type, then next *nρ*_2_ cells to the second type, etc.

Partitioned directed graphs are generated using SBM, with the vertices proportioned according to *ρ* (2), and then connected to other vertices with probabilities given by *P* (1). We label the vertices of *V* by *υ*_1_,…, *υ_n_*. Each directed graph is uniquely associated to an **adjacency matrix** *A*, an *n × n* binary matrix with the *ℓm*th entry given by 1 if (*υ_ℓ_, v_m_*) ∈ *E* and 0 otherwise. Note that since we consider only simple directed graphs in this work, the diagonal entries of the adjacency matrix are zero.

## 3 ADJACENCY SPECTRAL EMBEDDING

Given an *n × n* adjacency matrix *A* generated by a directed SBM, the goal is to predict the number of classes and recover the class assignment for each individual vertex of the graph, with no prior knowledge of *k, P* or *ρ*. The first step is to embed the adjacency matrix into a lower dimensional Euclidean space via singular value decomposition.

### 3.1 Singular Value Decomposition

Any real valued matrix *A* may be decomposed into a product *A = UDV^t^* where *D* is a diagonal matrix with nonnegative real entries, and *U* and *V* are real valued orthogonal matrices, called a **singular value decomposition**. We may choose *D* so that its entries, called the **singular values**, are nonnegative and weakly decreasing, in which case *D* is uniquely determined by A. The columns of U and V are called **singular vectors**.

In contrast, *U* and *V* are not unique; if the entires of *D* are distinct and nonzero, then *U* and *V* are determined up to a simultaneous factor of ±1 in each column of *U* and *V*. If there are repeating nonzero entries of *D*, the corresponding singular vectors span a subspace of dimension given by the number of copies of the repeated singular value. Any set of orthonormal vectors spanning this subspace can be used as the singular vectors in *U*, with a resulting choice in *V*. If any singular values vanish, the corresponding singular vectors in *U* and *V* may be chosen independently.

For any *d* ≤ rank(*A*), one can approximate *A* by a rank *d* decomposition

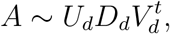

in which *U_d_* and *V_d_* are *n × d* matrices, and *D_d_* is a *d × d* diagonal matrix with nonnegative entries. Let 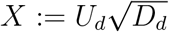 and 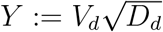, so that *A ~ XY^t^*.

### 3.2 Embedding in a Lower Dimension

We use a singular value decomposition of the adjacency matrix *A* to capture the most salient data in a low-dimensional space. We modify A by replacing the *i*th diagonal entry with the out-going degree of the *i*th vertex, divided by *n*−1. The out-going degree of the *i*th vertex is the number of out-going edges incident to the vertex, and is calculated by simply summing up all entries of the *i*th row of *A*. As *n* increases, this change in diagonal value has an insignificant impact on the matrix decomposition, assuming the graph is sparse. For each directed graph (V, E) and choice of embedding dimension *d*, the vectors forming the columns in the augmented matrix **X**: = [*X*|*Y*]^*t*^ provide a **dot product embedding** of A in a 2*d*-dimensional space. The columns of the concatenated matrix **X** are called **latent vectors**.

The optimal choice of *d* is a known open problem in literature, with no consensus on a best strategy. The necessity of selecting an optimum *d* is based on the fact that only a subset of the singular values of the high-dimensional data are informative and relevant to the subsequent statistical inference. Choosing a low *d* can result in discarding important information, while choosing a higher *d* than required not only increases computational cost but can adversely effect clustering performance due to the presence of extraneous variables which contribute towards noise in the data. For SBM graphs with large n, it has been shown (Fishkind et al. 2013) that the optimal choice of *d* is the rank of the block-connectivity matrix P, however in our context we assume no prior knowledge of P. A general methodology to choose the optimum value for *d* is then to examine the scree plot, the plot of the singular values in weakly decreasing order, and look for an ‘elbow-point’ which determines the cut-off between relevant and non-relevant dimensions based on the magnitude of the singular value. The scree plot for a SBM graph generated using the parameters of our surrogate model (1), (2), is shown in Fig. 1. Estimating the elbow-point using the unit invariant knee method (Christopoulos 2016) yields an optimum value of *d* = 4. This choice of *d* = 4 is also consistent if we instead use an alternative method (Satopaa et al. 2011) of estimating the distance from each point in the scree plot to a line joining the first and last points of the plot, and then selecting the elbow-point where this distance is the largest.

**Figure 1:**
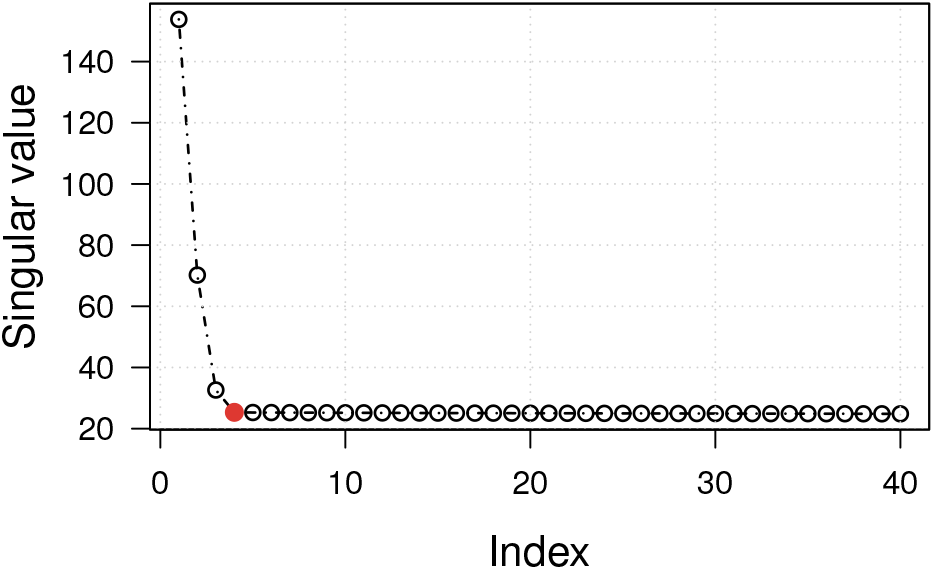
Model selection: *d* = 4 based on the elbow point of the scree plot of singular values (*n* = 16, 384). The top *d* singular values and their associated left- and right-singular vectors are concatenated to embed the graph in 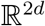.

We apply singular value decomposition directly to *A* before clustering, rather than to its Laplacian. For the case of a symmetric *A* (undirected graphs), under certain assumptions (Sussman et al. 2012), clustering of the resulting singular value decomposition converges to a negligible number of misclassified vertices. Such results have also been found in similar work applied to the Laplacian (Rohe et al. 2011; Vogelstein et al. 2019). However, to the best our knowledge, analogous results for directed graphs have not been explored.

## 4 GAUSSIAN MIXTURE MODEL BASED CLUSTERING

Let *A* be an *n × n* adjacency matrix and *A ~ XY^t^* be a singular decomposition with *d*-singular values. We denote by 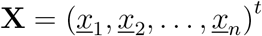 the data (latent vectors) obtained from this decomposition of *A*, where 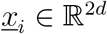 denotes the concatenation of *i*th row of *X* followed by the *i*th row of *Y*. Fig. 2 shows a scatterplot matrix of the latent vectors distributed in 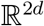, for the choice of embedding *d* = 4. The scatterplot depicts the data projected as points onto a two-dimensional subspace, whose coordinates are composed of a pair of the orthogonal singular vectors. The colors represent the original class assignment associated with each data point. The SBM graph was generated using the surrogate model (1), (2) for *k* = 8 classes, and *n* = 16, 384.

**Figure 2:**
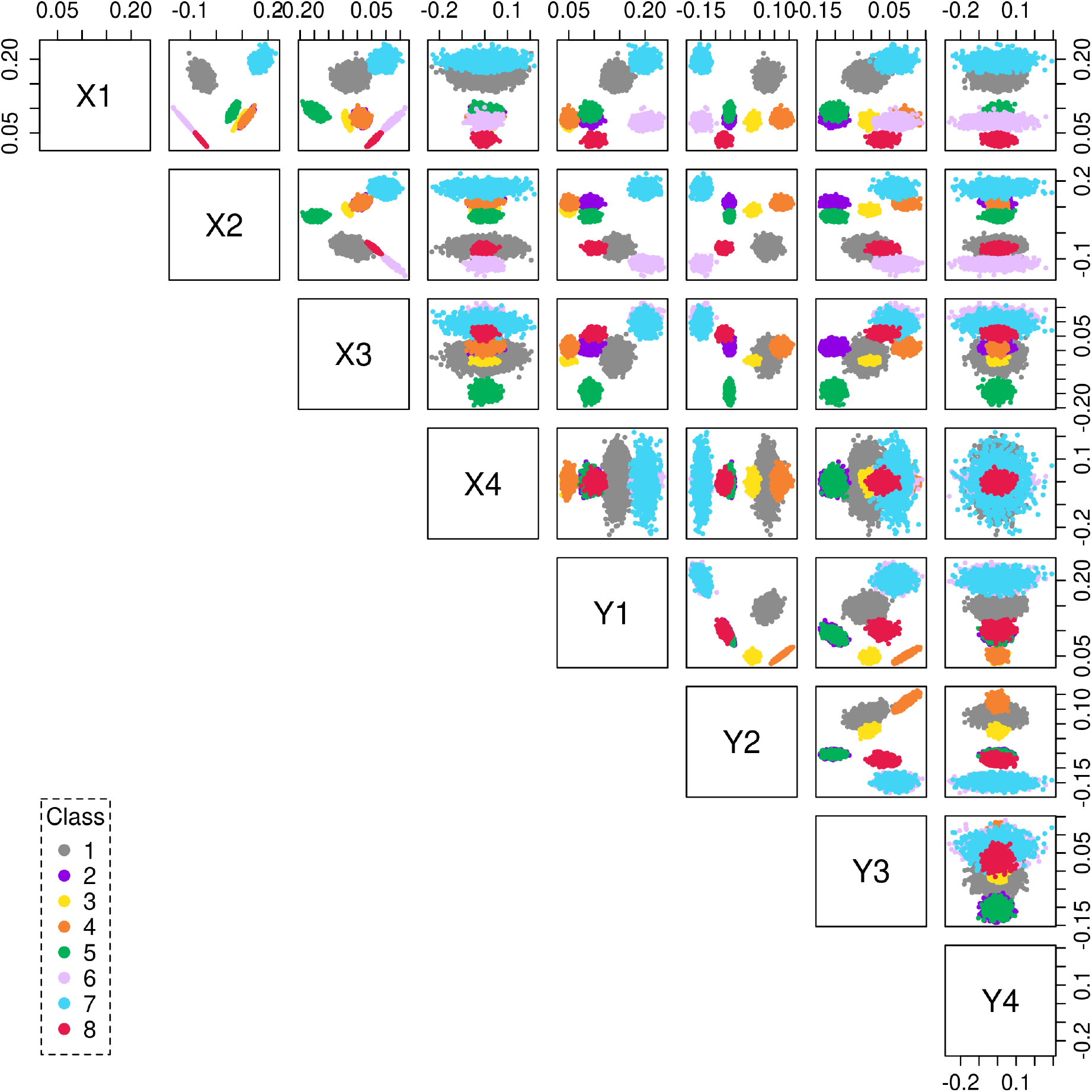
Scatterplot matrix showing the latent vectors of a SBM graph with *k* = 8 classes embedded in 2*d* = 8 dimensions. Each data point (*n* = 16, 384) is color coded as per its original class assigment.

### 4.1 Expectation-Maximization (EM) Algorithm

We cluster the data by modeling the latent vectors as a multivariate Gaussian mixture model (GMM) in order to predict the number of components, and the SBM block-partition function. For sufficiently dense graphs, and large *n*, the adjacency spectral embedding (ASE) central limit theorem demonstrates that *x_i_* behaves approximately as a random sample from a *k*-component GMM (Athreya et al. 2016).

Let 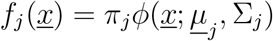, where 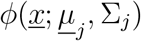 is the probability density function for the multivariate normal distribution with mean vector 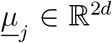, covariance matrix Σ_*j*_, and a component weight *π_j_* for *j* = 1,…, *κ*. The probability density function for the multivariate GMM with 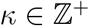 components is given by

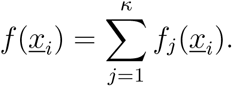

The Gaussian mixture model is fitted to the data using the expectation-maximization (EM) algorithm. We assume the Gaussian distributions may have aspherical covariances to address clusters in ellipsoidal shapes. The clusters are centered at the mean vector 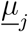, while other geometric features, such as the volume, shape and orientation, of each cluster are allowed to vary. Assuming the *n* data points 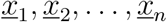 are independent draws,

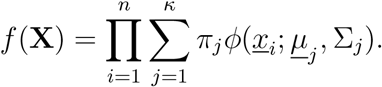

After an initialization of the mixture parameters 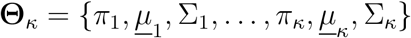, we set

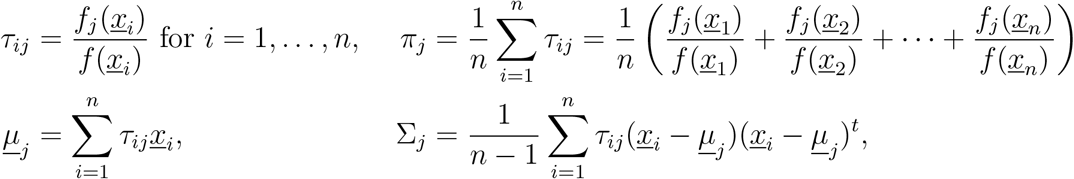

where the product 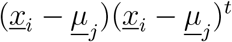 occurring in Σ_*j*_ is the tensor (outer) product.

The EM algorithm is used to iteratively improve upon the estimates by maximizing the log likelihood of the joint probability density function

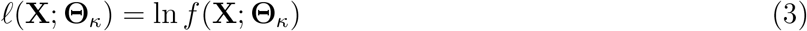

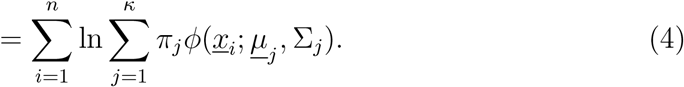

We iterate this process until convergence. After the first iteration, ∑_*j*_ *π_j_* = 1, and ∑_*j*_ *τ_ij_* = 1. This model assumes that 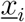 has an associated probability *τ_ij_* to be in each of the *j*th group. Indeed from this description we can define the estimated class assignment as follows: let *τ*: *V* → {1,…, *k*} be given by *τ*(*υ_i_*) = arg max_*j*_ *τ_ij_*.

### 4.2 Estimating the Number of Clusters

The model fitting procedure discussed above relies on a given number of GMM components *κ*, among which to distribute the *n* data points. Indeed, assigning each data point to its own cluster (*κ = n*) would uniquely identify connectivity behavior of each vertex, but would not illuminate common attributes. At the other extreme, *κ* = 1 provides no distinguishing information among vertices. Let *κ_min_* and *κ_max_* denote the smallest and largest values of practical interest for *κ*, respectively. We estimate the number of clusters by selecting the value of *κ* ∈ {*κ_min_*,…, *κ_max_*} which maximizes the Bayesian Information Criterion (BIC). BIC penalizes the model based on the number *p_κ_* of free parameters^2^, which grows linearly with *κ* and depends quadratically on the number of singular values *d*. Specifically, let 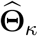 be the maximum likelihood estimate of the parameters given the data 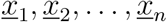 under the assumption that they are modeled by a multivariate Gaussian mixture model with *κ* components. The estimated number of classes is defined as

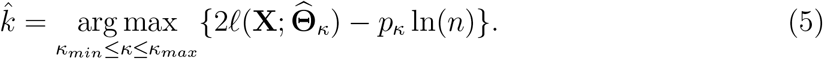

For each *κ*, the GMM fit results in a class assignment 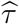 of each vector 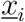 to a group labeled {1,…,*κ*}.

### 4.3 EM Initializations using Multiple Restarts

The final parameter estimates of the fitted model, are often sensitive to the initial values chosen to start the EM algorithm, especially for the case of finite mixture models (Shireman et al. 2017; Melnykov and Melnykov 2012). A poor initial choice of the model parameters may cause the EM algorithm to converge to a local but not global maximum of the likelihood function (Biernacki et al. 2003).

A workaround to the problem of EM initialization is the multiple restart approach (Biernacki et al. 2003; Kwedlo 2015). Specifically, given a set of data points, the EM algorithm is run *T* times (trials), each trial starting with different initial parameters. Each trial is run across all *κ* values with *κ_min_ ≤ κ ≤ κ_max_*, resulting in 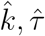 and a maximum BIC value for the trial. The final clustering is selected as the model with the highest BIC across all T trials. Considering the high prevalence of local maxima in the loglikelihood function, optimal solutions resulting from different trials are typically different. The highest BIC observed across a sufficiently large number of trials corresponds to the best estimate of the global maximum among local optima.

For each trial, an initial estimate of the model parameters is obtained by applying another preliminary clustering to the data. Towards this extent, we compare two variations of agglomerative hierarchical clustering. An inherent advantage of agglomerative hierarchical clustering is that it partitions the data simultaneously into any number of desired clusters, and that, for any trial, the initial clusters are similar across values of *κ*. In the first method, initial parameters are obtained by partitioning the data using random hierarchical agglomerative clustering (RHAC). In the second approach, initial parameters are obtained by applying model-based hierarchical agglomerative clustering (MBHAC) to a random subset of the data points. Both methods are described in further detail in the following subsections.

#### 4.3.1 Restarts using random hierarchical agglomerative clustering (RHAC)

At the outset RHAC begins with every data point 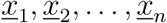 in its own cluster. Random pairs of clusters are then successively merged (with a uniform probability of choosing any two clusters for merging) until all *n* data points have been grouped into a single cluster. Equivalently, we could also start RHAC from a specific number of clusters, and successively proceed to form larger clusters. Since we do not know the true number of clusters we run EM for all values of 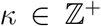, in the range *κ_min_ ≤ κ ≤ κ_max_*. Starting with an initial choice of *κ_max_* number of clusters, RHAC assigns each data point randomly to any one of the clusters, with uniform assignment probability 1/*κ_max_*. At the subsequent hierarchical agglomerative clustering stage, any two randomly picked clusters are combined, resulting in a total of *κ* − 1 clusters. This process is successively repeated until all data points have been grouped into *κ_min_* clusters. RHAC is computationally very efficient with a fast runtime, and a low memory usage cost of 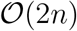.

During each trial we run the EM algorithm multiple (*κ_max_ − κ_min_* + 1) times on the data, incrementally decreasing the value of *κ* ∈ {*κ_min_,…, κ_max_*} each run. For each *κ*, the parameters of the randomly created RHAC partitions are used to start the EM. The EM algorithm is then run iteratively, maximizing the loglikelihood estimate, until convergence to an optimal solution. The proposed multiple restart RHAC based EM (mRHEM) algorithm is summarized in Algorithm 1.

##### Algorithm 1 *m*RHEM^†^

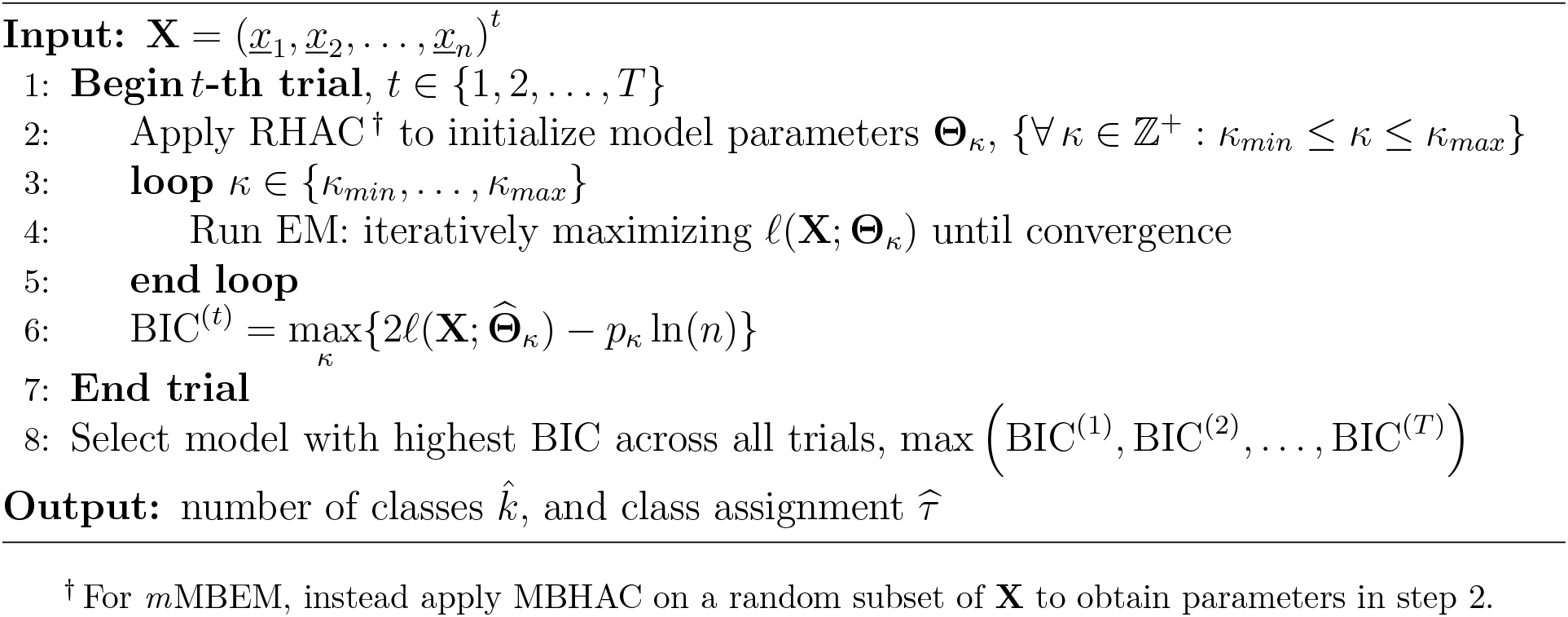

#### 4.3.2 Restarts using MBHAC on a random subset

Model-based hierarchical agglomerative clustering (MBHAC) uses a Gaussian mixturemodel to obtain a partition of the data (Fraley 1998; Scrucca and Raftery 2015), and is the default EM initialization method for the `mclust` R-package (Scrucca et al. 2016) Starting with each data point of the subset in its own cluster, MBHAC merges a pair of maximumlikelihood clusters at each successive stage of the hierarchical clustering, resulting in a partition for each *κ* ∈ {*n*,…, 1}. The parameters of these clusters obtained using MBHAC can then be used to initialize the EM algorithm across the desired range of *κ*.

Applying MBHAC to the full data set is deterministic, and computationally expensive with the memory usage cost being proportional to the square of the number of data points, 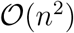 (Fraley 1998). As an alternate for large values of *n*, the initial model parameters can be obtained by applying MBHAC to a smaller subset of the data points chosen at random (with uniform probability) (Fraley 1998; Scrucca and Raftery 2015). The GMM is then fitted to all *n* data points by starting the EM algorithm with this choice of initial parameters.

We extend this randomized MBHAC approach to implement a multiple random restart version of the EM algorithm (*m*MBEM). Specifically, we run many trials on each data set. For each trial we choose a random subset from among the *n* data points and apply MBHAC to obtain the initial EM parameters for the desired range of mixture components *κ*. Finally, we select the model with the highest BIC across all trials. The *m*MBEM algorithm is therefore identical to *m*RHEM outlined in the previous section, with the only difference being the use of MBHAC applied to a random subset to initialize the model parameters (in step 2 of Algorithm 1).

### 4.4 The Probability Estimates

We obtain an estimate of the block connectivity probability matrix 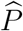 using the proportion of connected vertices given by our graph and using the partition 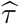. We define the *ij*th entry of this matrix by

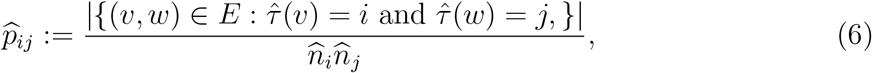

where 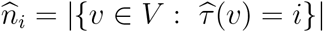. The ratio in (6) defines a value from 0 to 1.

The probability estimate is compared to the original parameters that generated the graph. Recall that *ρ_i_* is the proportion of vertices originally in the *i*th group, and *p_i_j* is the probability that a specified element of the *i*th group has a directed edge to a specified element in the *j*th group. The corresponding relative error rate is defined as

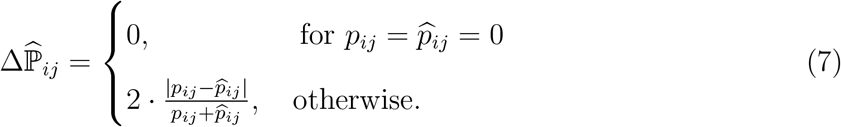

The percentage relative error in estimating the block-connection probabilities is a weighted average using the class proportions,

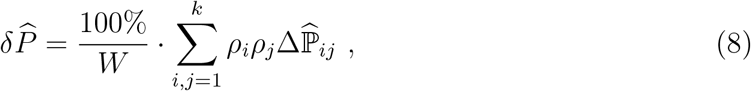

where 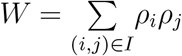, with the index set *I* = {(*i, j*): *p_ij_* ≠ 0, and 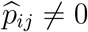}.

When the clustering is perfect, the expected difference 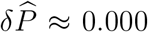 because perfect clustering implies that 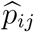 is the proportion of connected vertices in a size *n_i_n_j_* random sample from a binomial distribution with parameter *p_i_j*.

## 5 SIMULATION RESULTS

In order to validate the effectiveness of the proposed approach we performed multiple simulations using our surrogate connectome model. During the course of these simulations we randomly generated SBM graphs by systematically varying each of the parameters (*n, P, ρ*) of our surrogate model (1), (2). For each graph we performed ASE followed by GMM-based EM clustering. We compared the effects of EM initialization on clustering performance by applying the *m*RHEM and *m*MBEM algorithms, to the same graphs, respectively. Additionally, we also tested the robustness of our model to choices of embedding dimension *d*, the addition of noise, and the effect of varying the number of trials when applying multiple restart EM. We describe these results in detail below.

### 5.1 Varying the Embedding Dimension *d*

We first assess the impact that the choice of embedding dimension has on the clustering performance when using GMM-based hierarchical clustering. We generated 50 random graphs for each value of *n* using the surrogate model (1), (2), and then cluster them by embedding them in 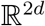 using ASE (varying the value of *d* each time).

For the sake of comparison, clustering was first performed by running the EM algorithm with initial parameters obtained by applying MBHAC to all *n* data points, implemented using the `mclust` R-package (Scrucca et al. 2016). Note that applying MBHAC to all data points creates deterministic partitions resulting in just a single trial, *T* = 1. Table 2 shows the percentage of 50 graphs in which the vertices were perfectly clustered (i.e., each vertex *υ_i_* was correctly assigned to its true class *τ*(*υ_i_*) by the algorithm) and the percentage of vertices that were misclassified across these graphs. The results indicate that using this approach to initialize the EM algorithm performed rather poorly, and was in general unsuccessful in clustering the latent vectors correctly. Interestingly, the method performed better for lower values of *d* and large *n*, with the misclassification rate being very low for these values.

**Table 2:** Clustering accuracy for EM initialization using MBHAC for a single trial, *T* = 1. The initial parameters were obtained by applying MBHAC to all *n* data points. *d* is the number of singular values chosen for ASE. A total of 50 graphs were used for each *n*.

| $n$ | % Perfect clustering (% vertices misclassified) | | | | |
| --- | --- | --- | --- | --- | --- |
| | $d = 2$ | $d = 3$ | $d = 4$ | $d = 5$ | $d = 6$ |
| $2^{12}$ (4, 096) | 2 (7.63) | 0 (14.55) | 2 (15.48) | 0 (18.18) | 0 (18.23) |
| $2^{13}$ (8, 192) | 14 (1.85) | 24 (7.45) | 12 (8.58) | 4 (10.83) | 2 (16.87) |
| $2^{14}$ (16, 384) | 28 (1.94) | 22 (2.76) | 12 (7.95) | 6 (8.11) | 2 (7.30) |
| $2^{15}$ (32, 768) | 34 (1.65) | 16 (2.76) | 12 (6.37) | 0 (5.26) | 0 (7.52) |

Tables 3 and 4 show the results when using the proposed multiple restart variations *m*MBEM, and *m*RHEM algorithms, respectively. Both algorithms were implemented with aid of the `mclust` package. A total of 100 trials were used to cluster each graph. We observe a drastic improvement in the clustering performance when using the random multiple restart approach. Also as expected, and in contrast to MBHAC, the clustering performance improves as *n* increases (Athreya et al. 2016).

**Table 3:** Clustering accuracy using *m*MBEM with T = 100 trials, wherein each trial was initialized using parameters obtained by applying MBHAC to a random subset of 2000 data points. A total of 50 graphs were used for each *n*.

| $n$ | % Perfect clustering (% vertices misclassified) | | | | |
| --- | --- | --- | --- | --- | --- |
| | $d = 2$ | $d = 3$ | $d = 4$ | $d = 5$ | $d = 6$ |
| $2^{12}$ (4, 096) | 0 (19.96) | 36 (6.10) | 14 (14.00) | 40 (0.04) | 44 (0.04) |
| $2^{13}$ (8, 192) | 0 (12.48) | 100 (0.00) | 58 (17.36) | 98 (0.01) | 98 (0.01) |
| $2^{14}$ (16, 384) | 14 (5.53) | 100 (0.00) | 78 (18.65) | 100 (0.00) | 100 (0.00) |
| $2^{15}$ (32, 768) | 100 (0.00) | 100 (0.00) | 26 (20.36) | 100 (0.00) | 100 (0.00) |

**Table 4:** Clustering accuracy using *m*RHEM with *T* = 100 trials. *d* is the number of singular values chosen for ASE. A total of 50 graphs were used for each *n*.

| $n$ | % Perfect clustering (% vertices misclassified) | | | | |
| --- | --- | --- | --- | --- | --- |
| | $d = 2$ | $d = 3$ | $d = 4$ | $d = 5$ | $d = 6$ |
| $2^{12}$ (4, 096) | 50 (0.03) | 46 (0.63) | 22 (5.05) | 10 (10.56) | 0 (9.72) |
| $2^{13}$ (8, 192) | 100 (0.00) | 100 (0.00) | 100 (0.00) | 98 (5.08) | 92 (7.17) |
| $2^{14}$ (16, 384) | 100 (0.00) | 100 (0.00) | 100 (0.00) | 100 (0.00) | 98 (3.31) |
| $2^{15}$ (32, 768) | 100 (0.00) | 100 (0.00) | 100 (0.00) | 100 (0.00) | 100 (0.00) |

For the results in Table 3, the size of the random subset used for *m*MBEM initialization was kept constant at 2000 data points, irrespective of the value of *n*. Rather surprisingly though, *m*MBEM performed poorly for the choice of embedding dimension d=4, which from Fig. 1 is the target dimension of interest. For the particular case of *d* = 4, we observed a consistent error pattern for all graphs that were not perfectly clustered. For these graphs the final clustering always resulted in 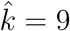, with the largest cluster being split into two.

The clustering results do improve when we increased the size of the random subset, but so does the computation time. In Table 5 we compare the performance of *m*MBEM as a function of the random subset size used for initialization, by applying it to the same 50 graphs each with *n* = 2^15^, and *d* = 4. The average CPU^3^ elapsed time shown is the time taken to perform agglomerative herirachial clustering given data X, and does not include the time taken to perform any other operation such as ASE, iterating EM, calculating the BICs, etc. Doubling the size of the random subset to 4000 data points led to approximately a six-fold increase in CPU computation time to perform randomized MBHAC, with only a marginal improvement in clustering accuracy. MBHAC initialization for subsets larger than 2000 points results in diminishing gain.

**Table 5:** Average time (in seconds) taken to perform different variations of agglomerative hierarchical clustering. A total of 50 graphs were used each with *n* = 2^15^, and *d* = 4.

| Intialization method | RHAC | MBHAC<br>(2000) | MBHAC<br>(4000) | MBHAC<br>(8000) | MBHAC<br>($2^{15}$)* |
| --- | --- | --- | --- | --- | --- |
| CPU Time (secs.) | 7.36 | 1.87 | 12.19 | 83.22 | 4554.31 |
| % Perfect clustering | 100 | 26 | 38 | 72 | 12* |

\* Applying MBHAC to all $n$ points, results in a single trial.

In contrast, *m*RHEM was largely insensitive to the choice of embedding dimensionality. It was also extremely consistent in its performance with near perfect clustering accuracy for *n* ≥ 2^13^. While we list results for 100 trials, a larger number of *m*RHEM trials resulted in even stronger results. Further, unlike *m*MBEM which is subject to an additional parameter (*υiz*. size of the random subset used for initialization) which directly affects its clustering accuracy and computational complexity, *m*RHEM is rather straightforward to implement and extremely efficient computationally. We use *m*RHEM exclusively for the remainder of the analysis.

### 5.2 Varying the Number of Vertices *n*

To examine the effects of varying *n* in further detail we fixed the choice of embedding dimensionality at a constant *d* = 4, as selected from Fig. 1. Table 6 shows the clustering performance of *m*RHEM with *T* = 100 trials for varying number of vertices. Misclassfied vertices were measured from maximal BIC among trials, and averaged over 50 graphs. Additionally, we also include the percentage relative error in estimating the block-connection probabilities (8), and measure the adjusted Rand index (ARI) (Hubert and Arabie 1985). Here the ARI was calculated in comparison to the true class memberships, and serves as an estimate for the overall accuracy of classification. ARI is a popular similarity score for comparing two partitioning schemes for the same data points, with a higher value of ARI indicating high similarity; 1 indicating that they are identical; and 0 for randomly generated partitions.

**Table 6:**
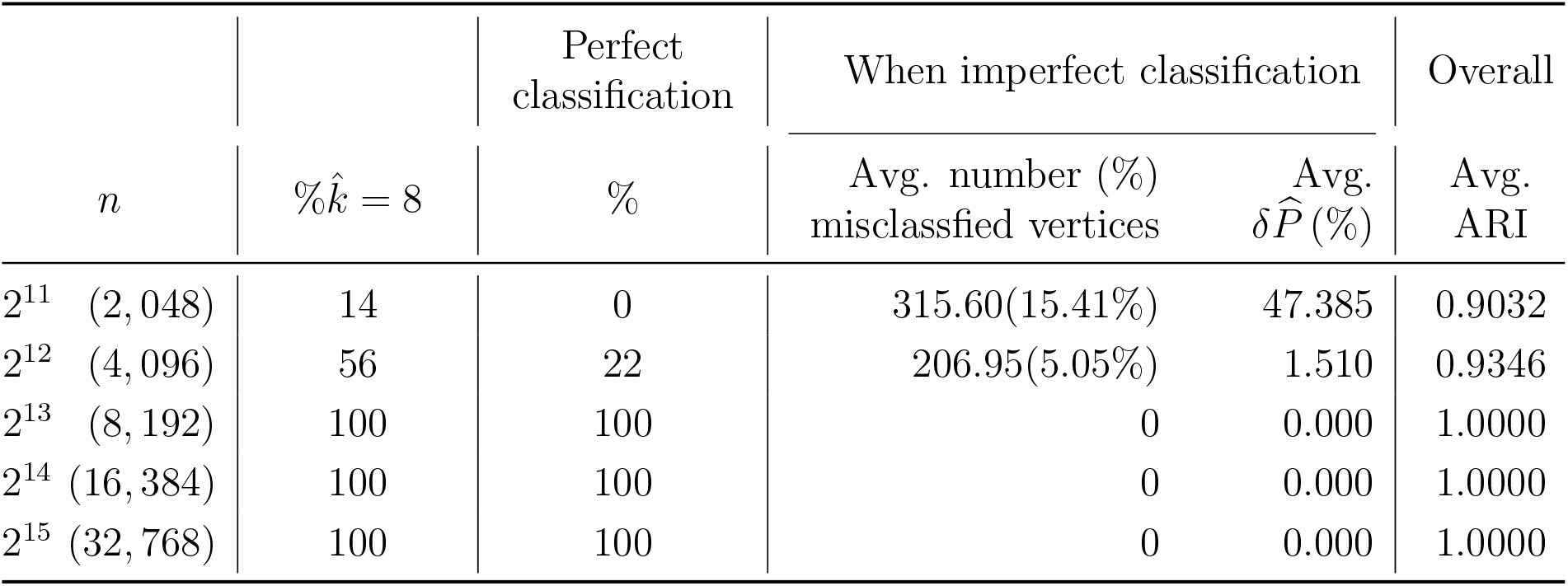
**Varying *n*:** Clustering accuracy using *m*RHEM with *T* = 100 trials for *d* = 4, as the number of vertices *n* is increased while keeping other parameters constant. A total of 50 graphs were used for each *n*.

|  |  | Perfect<br>classification | When imperfect classification |  | Overall |
| --- | --- | --- | --- | --- | --- |
| $n$ | $\% \hat{k} = 8$ | $\%$ | Avg. number (%)<br>misclassified vertices | Avg.<br>$\delta \hat{P} (\%)$ | Avg.<br>ARI |
| $2^{11}$ (2,048) | 14 | 0 | 315.60(15.41%) | 47.385 | 0.9032 |
| $2^{12}$ (4,096) | 56 | 22 | 206.95(5.05%) | 1.510 | 0.9346 |
| $2^{13}$ (8,192) | 100 | 100 | 0 | 0.000 | 1.0000 |
| $2^{14}$ (16,384) | 100 | 100 | 0 | 0.000 | 1.0000 |
| $2^{15}$ (32,768) | 100 | 100 | 0 | 0.000 | 1.0000 |

### 5.3 Varying the Proportions *ρ*

To test the robustness of the approach, we varied the SBM parameters, such that first *ρ* = (*ρ*_1_,…, *ρ_k_*) was varied while keeping *P* constant, and then *P* was varied while keeping *ρ* constant. To vary the class proportions we used a Dirichlet distribution *Dir*(*r_ρ_·ρ* + *J*_1;*k*_), where *r_ρ_* is a constant, and *J_i,j_* is an *i × j* matrix of all-ones. When *r_ρ_* = ∞ we have the original membership proportions in (2), and when *r_ρ_* = 0 the proportions are sampled from a uniform distribution. Table 7 shows the clustering results using *m*RHEM with 100 trials as *ρ* was varied. A total of 50 graphs were generated for each *ρ*, while keeping *P, n* = 2^14^, and *d* = 4 constant for each graph. We include the data for *r* = ∞ for comparison.

**Table 7:** **Varying *ρ*:** Clustering accuracy using *m*RHEM with 50 graphs and T = 100 trials, with varied block-membership proportions. Total number of vertices was kept constant *n* = 2^14^, and *d* = 4.

|  |  | Perfect<br>classification | When imperfect classification |  | Overall |
| --- | --- | --- | --- | --- | --- |
| $r_\rho$ | $\% \hat{k} = 8$ | % | Avg. number (%)<br>misclassified vertices | Avg.<br>$\delta \hat{P}$ (%) | Avg.<br>ARI |
| ∞ | 100 | 100 | 0 | 0.000 | 1.0000 |
| 10,000 | 100 | 100 | 0 | 0.000 | 1.0000 |
| 1,000 | 100 | 100 | 0 | 0.000 | 1.0000 |
| 100 | 100 | 100 | 0 | 0.000 | 1.0000 |
| 10 | 100 | 100 | 0 | 0.000 | 1.0000 |
| 0 | 78 | 78 | 3026.455(18.47%) | 2.743 | 0.9960 |

### 5.4 Varying the Probability Matrix *P*

To vary the connectivity probability matrix we used another Dirichlet distribution centered on *P*, with parameter *r*_p_, such that the probabilities are sampled from a uniform distribution when *r*_p_ = 0, and is given by the matrix *P* when *r*_p_ = ∞. Additionally, to ensure that the sampled graphs remain sparse we put bounds on the Dirichlet sampled *ij*-th entry of the probability matrix, 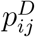, such that

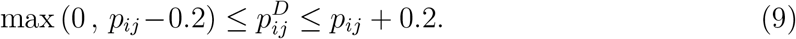

Table 8 shows the clustering results for varying *P* while keeping *ρ* constant for *n* = 4, 096. Alternatively, when the number of vertices is increased to *n* = 8,192, we observed that the *m*RHEM performance did not essentially deteriorate as block-connection probabilities were varied relative to the original values; when the number of vertices is set to *n* = 16, 384, *m*RHEM achieves perfect classification over the entire range of *r*_p_.

**Table 8:** **Varying *P*:** Clustering accuracy for using *m*RHEM with 50 graphs and T = 100 trials, with varied block-connection probabilities. Total number of vertices was kept constant *n* = 2^12^, and *d* = 4.

|  |  | Perfect<br>classification | When imperfect classification |  | Overall |
| --- | --- | --- | --- | --- | --- |
| $r_p$ | $\% \hat{k} = 8$ | % | Avg. number (%)<br>misclassified vertices | Avg.<br>$\delta \hat{P}$ (%) | Avg.<br>ARI |
| ∞ | 56 | 22 | 206.95(5.05%) | 1.510 | 0.9346 |
| 10,000 | 44 | 20 | 252.35(6.16%) | 11.184 | 0.9116 |
| 1,000 | 60 | 14 | 162.05(3.96%) | 11.000 | 0.9636 |
| 100 | 84 | 32 | 62.71(1.53%) | 6.754 | 0.9954 |
| 10 | 100 | 100 | 0 | 0.000 | 1.0000 |
| 0 | 100 | 100 | 0 | 0.000 | 1.0000 |

### 5.5 Effect of Adding Noise

To test the tolerance of the proposed clustering algorithm under experimentally realistic model misspecification we simulate errors in pre- or post-synaptic neuron identification. In order to do this we add noise to our model by randomly moving edges within the adjacency matrix. Specifically, a directed edge in the adjacency matrix is moved by flipping the corresponding 1 into a 0, and simultaneously flipping a randomly chosen 0 somewhere else in the matrix into a 1. Therefore, the total number of edges before and after the addition of noise remains the same while a subset of the edges is randomly (uniformly) scattered across all rows and columns of the matrix.

We measured how well *m*RHEM with *T* = 100 trials was able to estimate the original class assignment given a noisy graph. Fig. 3 shows the classification accuracy as a function of the proportion of edges moved. A total of 10 graphs were used each with *n* = 2^14^, and *d* = 4. The clustering results demonstrate *m*RHEM to be extremely tolerant towards added noise, with near prefect classification even with 50% edge misspecification.

**Figure 3:**
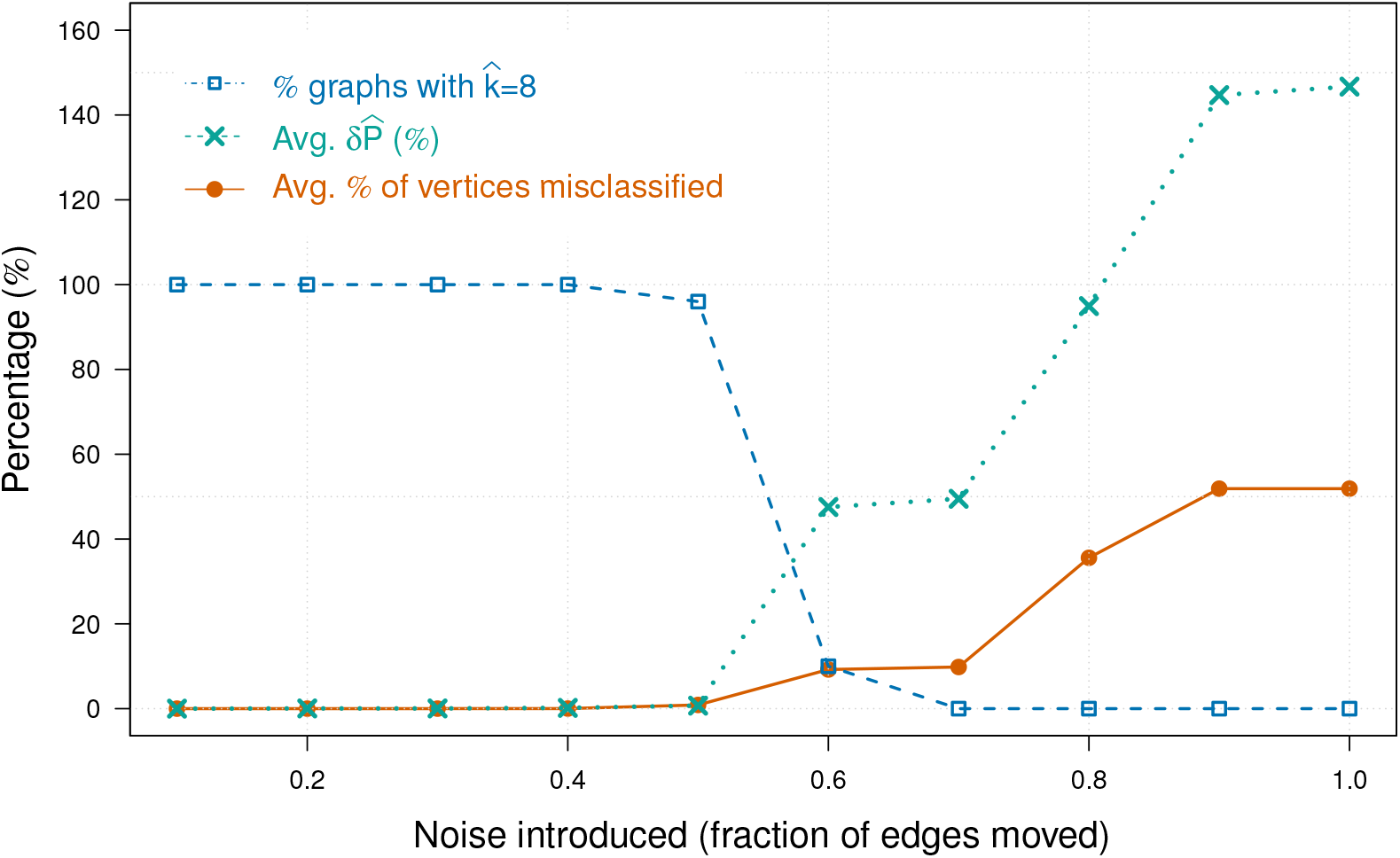
**Adding noise:** Average fraction of vertices that were misclassified versus the amount of noise added, i.e., number of moved edges. Also shown are the percentage of graphs whose clustering resulted in correctly estimating 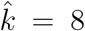, and relative error 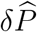, averaged over all graphs.

### 5.6 Influence of Number of Trials on *m*RHEM Performance

A fundamental disadvantage of using multiple restart EM is the computational cost associated with running multiple trials. To the best of our knowledge there is no theoretical solution available in the literature to determine the number of random initializations that would be sufficient to ensure a full examination of the likelihood function (Shireman et al. 2017; Biernacki et al. 2003). In the absence of an analytical solution, we use a numerical analysis to help determine the number of trials needed for *m*RHEM to converge to an optimal solution. Fig. 4 shows the percentage of graphs that are perfectly clustered as a function of the number of trials used to run *m*RHEM. Over 95% of the graphs for *r_ρ_* = ∞, and r*ρ* = 100 were perfectly clustered with only 37 trials.

**Figure 4:**
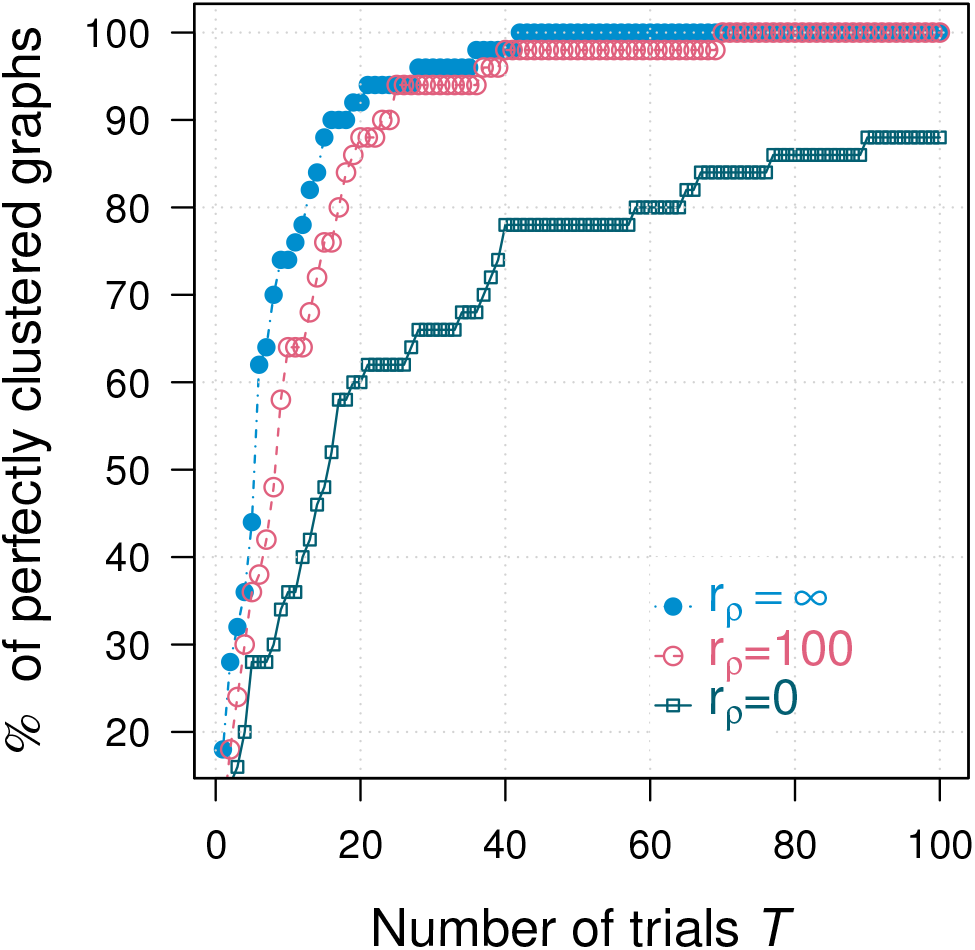
**Varying number of trials:** Percentage of perfectly clustered graphs when running T trials of *m*RHEM. A total of 50 graphs were used for different values of *r_ρ_* (with *r*_p_ = ∞, *n* = 2^14^, and *d* = 4 held constant).

Let BIC_*M*_ denote the BIC value for the optimal solution obtained when the EM is initialized by applying MBHAC to all *n* data points. For a single randomly chosen graph generated with the original parameters, Fig. 5(a) compares the number of misclassified vertices resulting from clustering when initializing using MBHAC, versus *m*RHEM with 100 trials, on the same graph. Trials are sorted along the horizontal axis to have increasing BIC values. While BIC_*M*_ is greater (has a better model fit) than ≈80% of the BIC^(*t*)^’s (data not shown), its ability to successfully predict class assignment is worse than ≈90% of the *m*RHEM trials, evidenced by the small number of data points among the *m*RHEM trials above the horizontal line indicating the number of misclassified vertices when initializing using MBHAC. A similar comparison is done for a single graph generated with *r*_ρ_ = 100 (Fig. 5 (b)) and for a single graph generated with *r_ρ_* = 0 (Fig. 5 (c)).

**Figure 5:**
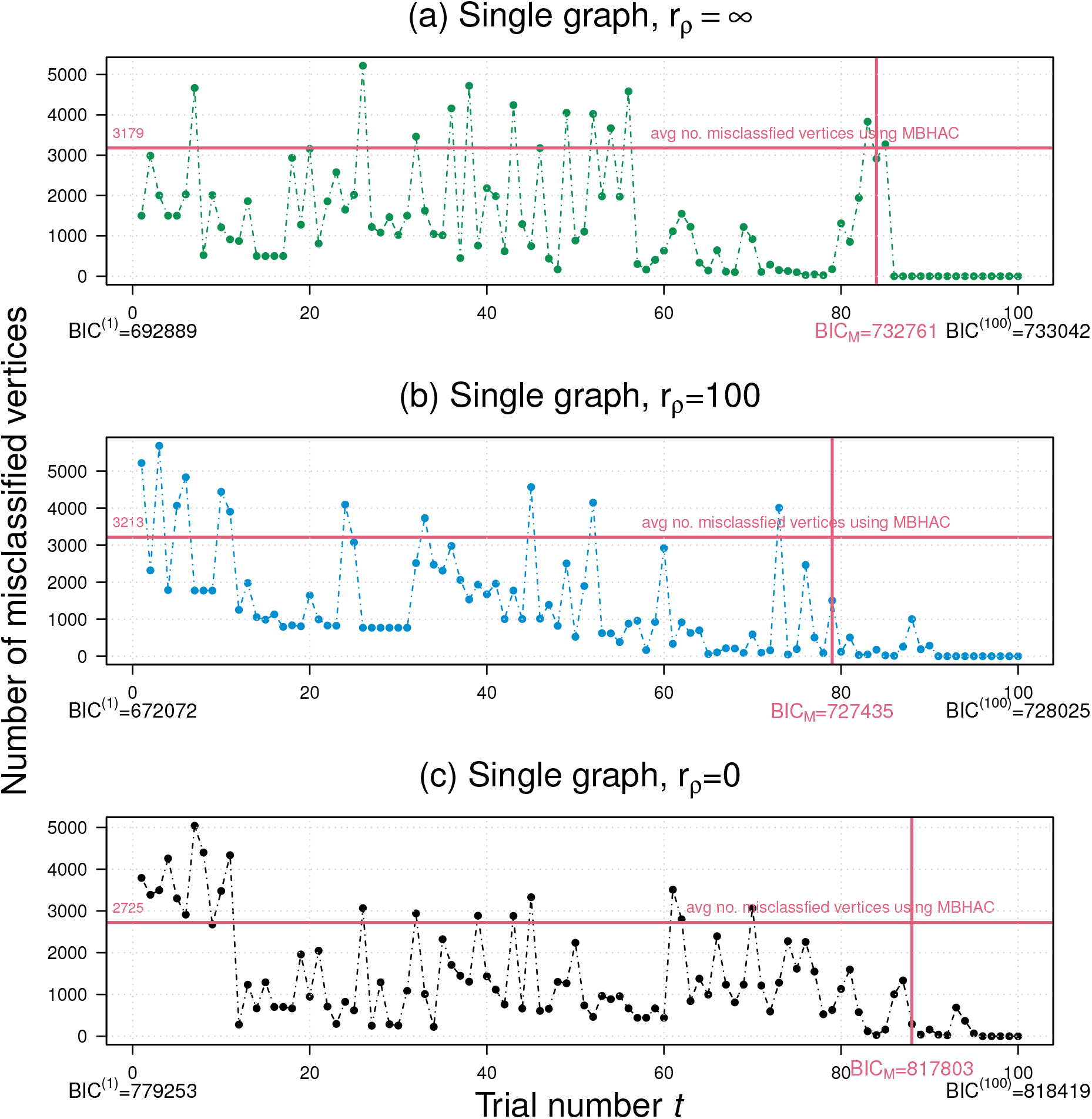
Number of misclassified vertices versus *m*RHEM trial number for a single graph for (a) *r_ρ_* = ∞, (b) *r_ρ_* = 100, and (c) *r_ρ_* = 0 (with *r*_p_ = ∞, *n* = 2^14^, and *d* = 4 held constant). The trials are sorted in increasing magnitude of BIC^(*t*)^. Also, shown for comparison is BIC_*M*_ corresponding to initialization using MBHAC applied to all *n* data points.

Despite the added computational cost associated with running EM several times, 100 *m*RHEM trials entails only a contained (≈ 270% on average) increase in CPU computation time. Additionally, since *m*RHEM is performing multiple quick trials, it allows for a relatively easy parallel-processing implementation (as opposed to one intensive trial using MBHAC). This could allow future CPU-intensive calculations to be performed simultaneously, resulting in significant time savings for *m*RHEM.

## 6 DISCUSSION

Understanding the types of neurons that comprise nervous systems is a fundamental step towards a more comprehensive understanding of neural circuits (Armañanzas and Ascoli 2015). The need for cell type classification from brain data is demonstrated by it being the first high-priority research areas identified by the Brain Research through Advancing Innovative Neurotechnologies (BRAIN) Initiative working group interim report^4^ and the resulting launch of the BRAIN Initiative Cell Census Network^5^. Previous approaches to classifying cell types have largely focused on the analysis of morphological, physiological or genetic properties. Here, we promote a complementary strategy that directly leverages connectivity. Our methodology effectively recovered the true number of clusters and cluster assignments as the number of vertices increased, even under experimentally realistic model misspecifications, corroborating its potential utility for analyzing real connectomic data.

Our ability as a community to estimate connectomes from real brain data has recently been transformed by breathtaking advances in techniques such as nanoscale electron microscopy (Denk and Horstmann 2004; Bock et al. 2011; Jarrell et al. 2012; Takemura et al. 2013), structural multi-color microscale light microscopy (Livet et al. 2007) paired with tissue clearing (Chung and Deisseroth 2013), functional mesoscale light microscopy (Ahrens et al. 2013; Schrödel et al. 2013), macroscale functional and diffusion magnetic resonance imaging (Craddock et al. 2013), and optical coherence tomography (Magnain et al. 2014). These technological breakthroughs require new approaches to analyze the resulting data, at scale, using principled statistical tools.

Our work illustrates the value of graph theoretic tools for discovering and assigning cell types in large scale simulations using connectivity information alone. In particular, we show that these methods can be used to recover class assignment for neural cells connected in biologically plausible proportions, at practical graph sizes for which data are emerging. The analysis and results of these surrogate data suggest that, at least in some circumstances, applying SVD and clustering techniques to the adjacency matrix rather than to its Laplacian results in consistent outcomes. However, there is a clear need for a theoretical framework that guarantees convergence for data that are asymmetric adjacency matrices representing directed graphs.

For GMM-based EM clustering of the surrogate data, the proposed *m*RHEM approach heavily outperforms the default MBHAC initialization used by `mclust` (Scrucca and Raftery 2015; Scrucca et al. 2016). We show that initializing the EM algorithm with random hierarchical agglomerative clustering multiple times is more effective than standard model-based hierarchical clustering at identifying the correct classification, as quantified by key measures of accuracy, such as clustering into the correct number of groups and misclassifying as few vertices as possible.

In future work, we will extend these results both theoretically and methodologically. We hope to characterize the circumstances in which one could expect better performance by *m*RHEM compared to MBAHC, and in particular find a probabilistic characterization of the optimal number of trials needed to obtain perfect clustering. While the proposed approach scales extremely well for large networks with 2^12^ ≤ *n* ≤ 2^15^ vertices, in order to model networks with even wider range and complexity, it is necessary to further examine the relationship between the parameter k and the required *n* for effective clustering. Such assessments could drive experimental efforts to reach benchmarked data collection goals. We also hope to extend these results to include not only connectivity information, but also various vertex and edge attributes, such as spatial, morphological, electrophysiological, and genetic properties. Finally, we strive to apply these methods to estimated connectomes from biological data to foster novel neuroscientific insights.

## SUPPLEMENTARY MATERIAL

**R code:** To generate surrogate data, and replicate simulation results described in the article. (`GNU zipped tar file`)

## Footnotes

1 These are called simply SBMs in some cases, such as Sussman et al. (2012).

2 More specifically, 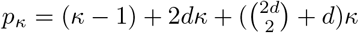.

3 Desktop workstation with eight-core AMD Ryzen 2700x (3.7GHz) with 32GB RAM (DDR4, 3200MHz), and `mclust` version 5.4.2.

4 http://www.nih.gov/science/brain/11252013-Interim-Report-ExecSumm.pdf

5 http://biccn.org

## REFERENCES

Abbe, E. (2018), “Community detection and stochastic block models: Recent developments,” Journal of Machine Learning Research, 18, 1–86. 5

Ahrens, M. B., Orger, M. B., Robson, D. N., Li, J. M., and Keller, P. J. (2013), “Whole-brain functional imaging at cellular resolution using light-sheet microscopy,” Nature Methods, 10, 413. 27

Armañanzas, R. and Ascoli, G. A. (2015), “Towards the automatic classification of neurons,” Trends in Neurosciences, 38, 307–318. 27

Ascoli, G. A., Alonso-Nanclares, L., Anderson, S. A., Barrionuevo, G., Benavides-Piccione, R., Burkhalter, A., Buzsáki, G., Cauli, B., DeFelipe, J., Fairén, A., et al. (2008), “Petilla terminology: nomenclature of features of GABAergic interneurons of the cerebral cortex,” Nature Reviews Neuroscience, 9, 557. 2

Athreya, A., Priebe, C. E., Tang, M., Lyzinski, V., Marchette, D. J., and Sussman, D. L. (2016), “A limit theorem for scaled eigenvectors of random dot product graphs,” Sankhya A, 78, 1–18. 12, 19

Betzel, R. F., Medaglia, J. D., and Bassett, D. S. (2018), “Diversity of meso-scale architecture in human and non-human connectomes,” Nature Communications, 9, 1–14. 3

Biernacki, C., Celeux, G., and Govaert, G. (2003), “Choosing starting values for the EM algorithm for getting the highest likelihood in multivariate Gaussian mixture models,” Computational Statistics & Data Analysis, 41, 561–575. 3, 14, 24

Bock, D. D., Lee, W.-C. A., Kerlin, A. M., Andermann, M. L., Hood, G., Wetzel, A. W., Yurgenson, S., Soucy, E. R., Kim, H. S., and Reid, R. C. (2011), “Network anatomy and in vivo physiology of visual cortical neurons,” Nature, 471, 177. 27

Christopoulos, D. (2016), “Introducing Unit Invariant Knee (UIK) as an objective choice for elbow point in multivariate data analysis techniques,” Available at SSRN 3013076. 9

Chung, K. and Deisseroth, K. (2013), “CLARITY for mapping the nervous system,” Nature Methods, 10, 508. 27

Craddock, R. C., Jbabdi, S., Yan, C.-G., Vogelstein, J. T., Castellanos, F. X., Di Martino, A., Kelly, C., Heberlein, K., Colcombe, S., and Milham, M. P. (2013), “Imaging human connectomes at the macroscale,” Nature Methods, 10, 524–539. 27

DeFelipe, J., López-Cruz, P. L., Benavides-Piccione, R., Bielza, C., Larrañaga, P., Anderson, S., Burkhalter, A., Cauli, B., Fairén, A., Feldmeyer, D., et al. (2013), “New insights into the classification and nomenclature of cortical GABAergic interneurons,” Nature Reviews Neuroscience, 14, 202. 2

Denk, W. and Horstmann, H. (2004), “Serial block-face scanning electron microscopy to reconstruct three-dimensional tissue nanostructure,” PLOS Biology, 2, e329. 27

Faskowitz, J., Yan, X., Zuo, X.-N., and Sporns, O. (2018), “Weighted stochastic block models of the human connectome across the life span,” Scientific Reports, 8, 1–16. 3

Fishkind, D. E., Sussman, D. L., Tang, M., Vogelstein, J. T., and Priebe, C. E. (2013), “Consistent adjacency-spectral partitioning for the stochastic block model when the model parameters are unknown,” SIAM Journal on Matrix Analysis and Applications, 34, 23–39. 9

Fraley, C. (1998), “Algorithms for model-based Gaussian hierarchical clustering,” SIAM Journal on Scientific Computing, 20, 270–281. 15, 16

Holland, P. W., Laskey, K. B., and Leinhardt, S. (1983), “Stochastic blockmodels: First steps,” Social Networks, 5, 109–137. 5

Holland, P. W. and Leinhardt, S. (1981), “An exponential family of probability distributions for directed graphs,” Journal of the american Statistical association, 76, 33–50. 5

Hubert, L. and Arabie, P. (1985), “Comparing partitions,” Journal of Classification, 2, 193–218. 21

Jarrell, T. A., Wang, Y., Bloniarz, A. E., Brittin, C. A., Xu, M., Thomson, J. N., Albertson, D. G., Hall, D. H., and Emmons, S. W. (2012), “The connectome of a decision-making neural network,” Science, 337, 437–444. 27

Kwedlo, W. (2015), “A new random approach for initialization of the multiple restart EM algorithm for Gaussian model-based clustering,” Pattern Analysis and Applications, 18, 757–770. 3, 14

Livet, J., Weissman, T. A., Kang, H., Draft, R. W., Lu, J., Bennis, R. A., Sanes, J. R., and Lichtman, J. W. (2007), “Transgenic strategies for combinatorial expression of fluorescent proteins in the nervous system,” Nature, 450, 56. 27

Magnain, C., Augustinack, J. C., Reuter, M., Wachinger, C., Frosch, M. P., Ragan, T., Akkin, T., Wedeen, V. J., Boas, D. A., and Fischl, B. (2014), “Blockface histology with optical coherence tomography: a comparison with Nissl staining,” NeuroImage, 84, 524–533. 27

Melnykov, V. and Melnykov, I. (2012), “Initializing the EM algorithm in Gaussian mixture models with an unknown number of components,” Computational Statistics & Data Analysis, 56, 1381–1395. 14

Moyer, D., Gutman, B., Prasad, G., Faskowitz, J., Steeg, G. V., and Thompson, P. (2015), “Blockmodels for connectome analysis,” in 11th International Symposium on Medical Information Processing and Analysis, International Society for Optics and Photonics, Cuenca, Ecuador: SPIE, vol. 9681, pp. 62–70. 3

Pavlovic, D. M., Vértes, P. E., Bullmore, E. T., Schafer, W. R., and Nichols, T. E. (2014), “Stochastic blockmodeling of the modules and core of the Caenorhabditis elegans connectome,” PLOS One, 9, e97584. 3

Priebe, C. E., Park, Y., Tang, M., Athreya, A., Lyzinski, V., Vogelstein, J. T., Qin, Y., Cocanougher, B., Eichler, K., Zlatic, M., and Cardona, A. (2017), “Semiparametric spectral modeling of the Drosophila connectome,” Preprint arXiv:1705.03297. 3

Priebe, C. E., Park, Y., Vogelstein, J. T., Conroy, J. M., Lyzinski, V., Tang, M., Athreya, A., Cape, J., and Bridgeford, E. (2019), “On a two-truths phenomenon in spectral graph clustering,” Proceedings of the National Academy of Sciences, 116, 5995–6000. 3

Rohe, K., Chatterjee, S., Yu, B., et al. (2011), “Spectral clustering and the high-dimensional stochastic blockmodel,” The Annals of Statistics, 39, 1878–1915. 10

Satopaa, V., Albrecht, J., Irwin, D., and Raghavan, B. (2011), “Finding a “Kneedle” in a Haystack: Detecting knee points in system behavior,” in 31st International Conference on Distributed Computing Systems Workshops, Minneapolis, MN, USA, pp. 166–171. 9

Schrödel, T., Prevedel, R., Aumayr, K., Zimmer, M., and Vaziri, A. (2013), “Brain-wide 3D imaging of neuronal activity in Caenorhabditis elegans with sculpted light,” Nature Methods, 10, 1013. 27

Scrucca, L., Fop, M., Murphy, T. B., and Raftery, A. E. (2016), “mclust 5: clustering, classification and density estimation using Gaussian finite mixture models,” The R Journal, 8, 289–317. 16, 18, 28

Scrucca, L. and Raftery, A. E. (2015), “Improved initialisation of model-based clustering using Gaussian hierarchical partitions,” Advances in data analysis and classification, 9, 447–460. 4, 15, 16, 28

Shireman, E., Steinley, D., and Brusco, M. J. (2017), “Examining the effect of initialization strategies on the performance of Gaussian mixture modeling,” Behavior Research Methods, 49, 282–293. 3, 14, 24

Sussman, D. L., Tang, M., Fishkind, D. E., and Priebe, C. E. (2012), “A consistent adjacency spectral embedding for stochastic blockmodel graphs,” Journal of the American Statistical Association, 107, 1119–1128. 3, 5, 10

Takemura, S.-y., Bharioke, A., Lu, Z., Nern, A., Vitaladevuni, S., Rivlin, P. K., Katz, W. T., Olbris, D. J., Plaza, S. M., Winston, P., et al. (2013), “A visual motion detection circuit suggested by Drosophila connectomics,” Nature, 500, 175. 27

Vogelstein, J. T., Bridgeford, E. W., Pedigo, B. D., Chung, J., Levin, K., Mensh, B., and Priebe, C. E. (2019), “Connectal coding: Discovering the structures linking cognitive phenotypes to individual histories,” Current Opinion in Neurobiology, 55, 199–212. 10

Wheeler, D. W., White, C. M., Rees, C. L., Komendantov, A. O., Hamilton, D. J., and Ascoli, G. A. (2015), “Hippocampome.Org: A knowledge base of neuron types in the rodent hippocampus,” Elife, 4, e09960. 6

